# Back to Basics: Spectrum and Peptide Sequence are Sufficient for Top-tier Mass Spectrometry Proteomics Identification

**DOI:** 10.1101/2024.08.19.606805

**Authors:** Maximilien Burq, Dejan Stepec, Juan Restrepo, Jure Zbontar, Shamil Urazbakhtin, Bryan Crampton, Shivani Tiwary, Rehan Chinoy, Melissa Miao, Juergen Cox, Peter Cimermancic

## Abstract

The original mass spectrometry search engines used simple algorithms for peptide identification. Recent tools improved accuracy by adding several extra components such as fragment ion intensities or retention times prediction and training target-decoy classifiers on-the-fly, leading to sometimes inconsistent results.

Our study explores the impact of replacing those extra components with a deep-learning pretrained model that directly learns the complex relationship between the full spectra and associated peptide sequence, without using decoys. This simplified workflow has fewer parameters to tweak, making it easier to use and perform robustly on data from instruments and use-cases never seen during training.

Surprisingly, our approach consistently identifies more peptides than FragPipe, PEAKS, and Proteome Discoverer (12%, 9%, and 21% more, respectively, across a range of datasets). Tesorai Search is also fast – 250 immunopeptidomics searches in 45 minutes – and free for academics, available as a webserver at console.tesorai.com.

## Introduction

Mass spectrometry (MS)-based proteomics typically involves breaking down complex protein mixtures into smaller peptides, which are then ionized and measured by a mass spectrometer. The instrument records the mass-to-charge ratio (m/z) of these intact peptides and subsequently fragments them to generate tandem mass spectra (MS2). Database search engines are the cornerstone computational tools used to interpret these complex spectra to determine the original peptide sequences. They work by comparing the experimentally acquired spectra against theoretical spectra generated from known protein sequences stored in databases. This process typically involves filtering potential peptide candidates based on precursor mass and then scoring the match between the experimental fragment ions and the predicted fragments for each candidate, resulting in peptide-spectrum matches (PSMs). The accuracy and comprehensiveness of protein identification are critically dependent on the performance of these search engines.

Original database search engines for mass spectrometry proteomics, such as Comet [1], MaxQuant (Andromeda) [2], [3], MSFragger [4], MS-GF+ [5], Proteome Discoverer [6] and Sequest [7], rely solely on spectral data — using precursor m/z for candidate selection and employing simple barcode matching with fixed canonical fragment ion intensities for scoring peptide-spectrum matches (PSMs). More recent search engines and rescoring tools such as MSBooster [8], MS2Rescore [9], PEAKS [10], Sage [11], Prosit/Oktoberfest [12, 13], Inferys [14] or Chimerys [15], have dramatically improved peptide identification rates with scoring innovations that incorporated dozens to hundreds of hand-crafted features based on prediction of canonical fragment ion intensities, differences between predicted and measured retention time, peptide lengths, and amino acid composition, followed by on-the-fly training of target-decoy classifiers such as Percolator [16] or PeptideProphet [17]. This added complexity increases the overfitting, which can in turn lead to underestimating the true false-discovery rate (FDR) in the generated search outputs [18].

We postulate that the full potential of proteomics identification can be achieved by returning to a spectrum-only approach — one that fully exploits the rich information in the entire spectrum alongside the peptide sequence. We propose that the limitations of the traditional methods arose not from the inadequacy of spectral data per se, but rather from incomplete utilization of the spectrum’s information, insufficient incorporation of peptide sequence context, and reliance on scoring functions that fail to capture complex, non-linear associations between spectra and sequence. To test that hypothesis, we introduce an end-to-end deep-learning model designed to receive the complete spectrum and peptide sequence and directly output a calibrated score for a PSM. This model, scaled to capture chemical complexity and trained on heterogeneous datasets covering various enzymes, instruments, and fragmentation types, employs a novel loss function that avoids reliance on decoy training. This is especially important as models get larger and the risk that the model will learn entire proteomes increases. Our results demonstrate that a spectrum-based approach not only achieves high accuracy and sensitivity in peptide-spectrum matching but also surpasses the performance of both traditional and recent search engines. These results are in contrast to approaches in [19, 20, 21, 22] which share some of our ideas, but were unable to outperform modern tools for database search.

Achieving high performance without on-the-fly model retraining required several breakthroughs, both on the model architecture and training paradigm. Similarly to traditional search engines, we pre-compute theoretical fragment ion intensities for the candidate peptide sequence. The model, however, also has access to the full measured spectrum, including regions that are outside of those theoretical fragment m/z values. We propose a new modular architecture where the model propagates its own embedding vectors for each breakpoint, enabling it to implicitly learn features related to precursor fragmentation and fragment ion ionization probability. For training, we introduce a new paradigm where the model learns to discriminate correct PSMs from high-scoring incorrect ones. Thus, decoys are not used at any point during model training and are reserved for (FDR) estimation. This is critical to ensure that the model does not overfit to the decoy generation process, which would yield incorrect results.

Our secondary contribution is an extensive benchmarking of nine widely used database search engines, which we ran on diverse datasets, including tryptic digests, immunopeptidomics, and single-cell analyses. We find that Tesorai Search consistently identifies more unique peptides compared to established search engines like MaxQuant, FragPipe (+MSBooster), PEAKS, and Proteome Discoverer, achieving average increases of 12% to 68%. We hope that this benchmark can serve as the basis for accurate comparisons, spurring further innovation and enhancements of existing and new search engines.

Further experiments confirm the model’s robustness, showing strong performance even on data not encountered during training, such as data produced with different instrument types (TOF) or using isobaric labeling (TMT). We verify through entrapment analyses that these increased identifications are achieved while maintaining accurate FDR control. Ablation studies confirm the model effectively utilizes information about peptide sequences and fragmentation spectra, including peaks coming outside of the canonical fragment ion series. Finally, we present a scalable, user-friendly cloud implementation capable of processing large datasets rapidly, making this advanced model readily accessible to the research community.

## Results

### Pretrained model paradigm

We hypothesized that training a single large peptide-spectrum matching algorithm can increase peptide identifications across use cases and reduce search engine complexity and data processing steps. To test this hypothesis, we propose a new algorithmic paradigm in which a deep-learning model outputs a score directly from a single tandem mass spectrum and candidate peptide sequence. This contrasts with existing ML-powered tools, which compute dozens to hundreds of metrics. In some cases, such as ion intensity or retention time prediction, each metric requires its own deep-learning model. These metrics then need to be combined into a score using a second-stage ML model – typically Percolator or PeptideProphet. Furthermore, to boost the performance, Percolator-based approaches incorporate information unrelated to peptide-spectrum matching, like charge and retention time error. We show that these additional features are unnecessary for accurate peptide-spectrum matching.

The Tesorai Search workflow (Fig. 1A) starts with user-provided raw mass spectrometry data (.d, .raw, or .mzml) and a protein sequence database (FASTA). Candidate PSMs are generated using Comet, MaxQuant, and MSFragger with permissive settings (100% FDR). Our pre-trained deep-learning model then rescores each spectrum-sequence pair, outputting a single score. These scores are subsequently used in a standard target-decoy competition strategy (see Methods) for accurate FDR control. The system outputs high-confidence PSMs and peptide identifications via a scalable cloud infrastructure.

**Figure 1.**
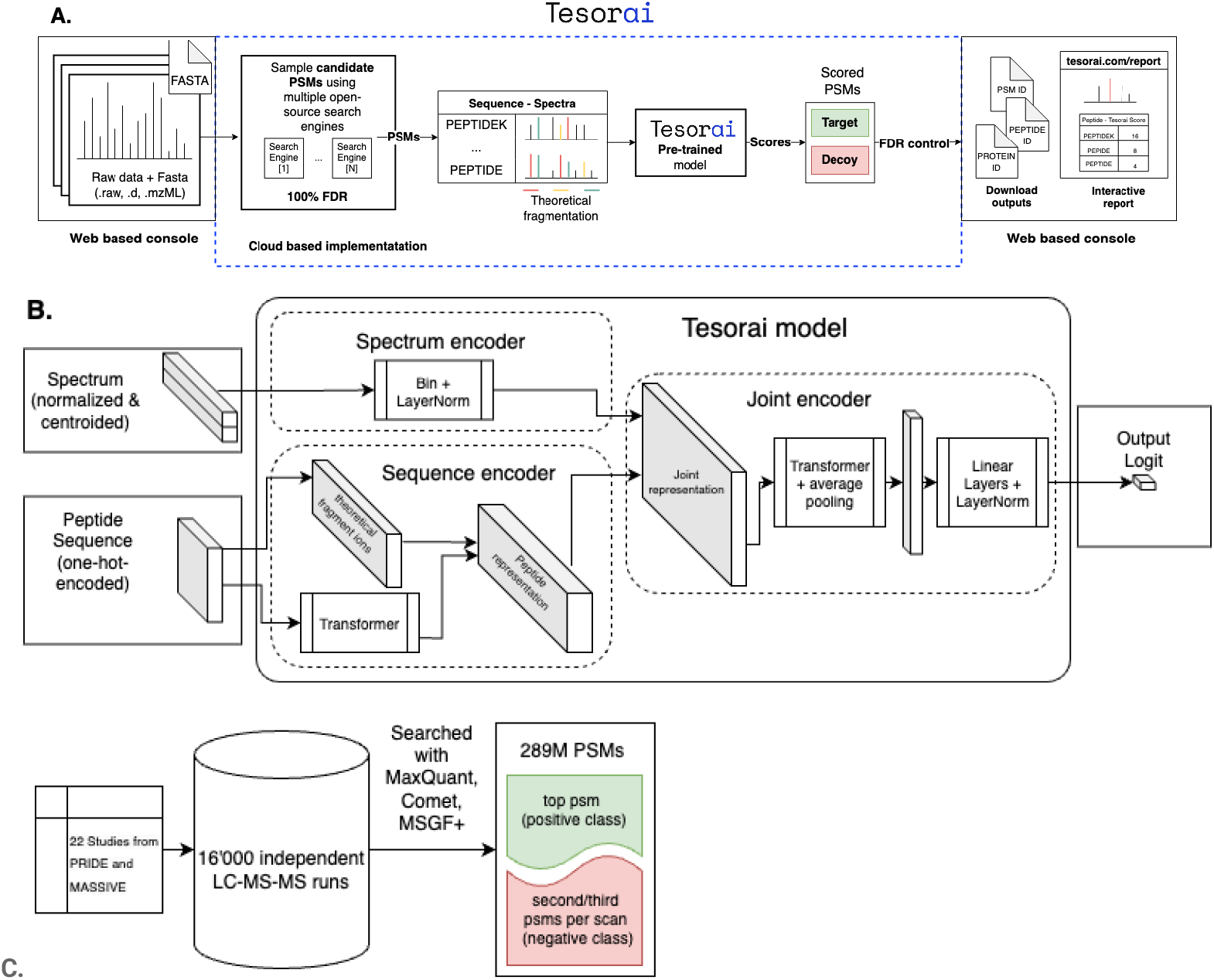
Overview of Tesorai Search. **A. Rescoring pipeline**. Illustrates the steps from raw data processing to rescoring with the pretrained model to the final list of identified peptides after FDR control. **B. Deep-learning-based rescoring model architecture**. Input data (MS2 spectrum and peptide sequence) is processed by the three key components, spectrum encoder, sequence encoder, and joint encoder, to generate a PSM score. **C. Training dataset generation and curation**. The training set consists of a pool of 22 studies and 16.000 LC-MS-MS runs processed with MaxQuant, Comet, and MS-GF+. The top PSM was taken as the positive class, and the second and third as the negative class.

Our approach simplifies processing by eliminating custom steps like deisotoping or recalibration, as these are implicitly learned by the model. The model architecture (Fig. 1B; Methods) has three components. A sequence encoder processes the modified peptide sequence (one-hot encoded), computes theoretical fragments, and combines them with a learned sequence embedding into a vector representation. A spectrum encoder generates a vector representation from the input centroided tandem mass spectrum (m/z and intensity lists). A joint encoder integrates the peptide and spectrum vectors, outputting a single numerical score indicating match quality.

We trained the model on a diverse dataset of ∼289 million tandem mass spectra from 16,000 runs across 22 studies (Fig. 1C; Methods). Spectra were searched using MaxQuant, MS-GF+, and Comet to identify top-ranking (positive examples) and lower-ranked (negative examples) peptide sequences. Importantly, the training occurs only once, does not utilize decoy sequences, and does not require retraining on the specific dataset of interest. Using real sequences in both positive and negative classes forces the model to learn genuine sequence-spectrum associations rather than memorize peptidomes.

### Increased number of identifications across many common use cases

We benchmarked Tesorai Search against leading search engines: MaxQuant [2], FragPipe (with MSBooster [7]), Proteome Discoverer (using Chimerys/Inferys [17] — see Methods), and PEAKS [9] (Fig. 2). All tools, including Tesorai Search, were run with default settings on seven diverse datasets representing common proteomics use cases. To ensure fairness, some datasets were chosen from the original benchmark publications of the compared engines. Our open-source data-processing code is available on GitHub^1^.

**Figure 2:**
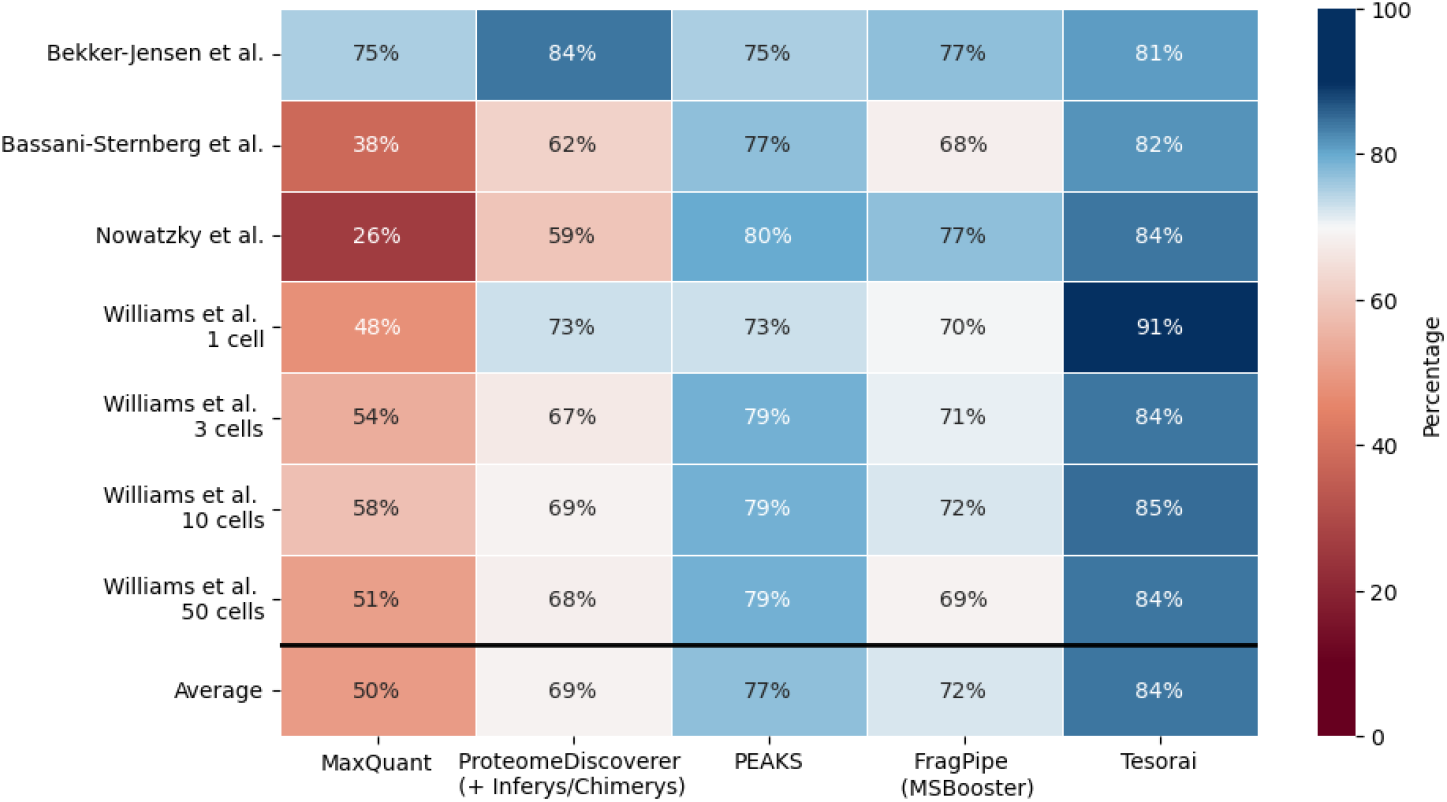
Unique peptide identifications (without modifications) at 1% FDR. Numbers are shown relative to the union of all identified peptides across all search engines. **Bekker-Jensen**. Deep fractionated tryptic HeLa sample **[23]. Bassani-Sternberg** Immunopeptidomics use-case **[24]. Nowatzky**. Immunopeptidomics use-case; MSV000089312 **[25]. Williams** Single-cell samples (NanoPots) **[26]**. The bottom row shows the average across all samples.

We selected a deeply fractionated tryptic HeLa sample from Bekker-Jensen et al. [23], which was previously analyzed in both Prosit and MSBooster publications. This sample was searched against a human FASTA file obtained from the original Prosit publication. Subsequently, two immunopeptidomics samples were chosen. The first originated from Bassani-Sternberg et al. [24], where a sample designated Mel15 had been utilized in benchmarking Prosit and MSBooster. The second dataset was for Behçet’s disease (Cavers et al.) [25]. Lastly, to assess our model’s performance on samples with limited quantities, we incorporated a study encompassing single-cell and few-cell analysis [26], previously used to assess performance in the MSBooster publication.

On average, we identified 17% more (non-modified) peptides than FragPipe/MSBooster, 9% more than PEAKS, 21% more than Proteome Discoverer (+ Chimerys/Inferys), and 68% more than MaxQuant. In Figure 2, we list the number of peptides identified by each search engine, normalized as a fraction of the total number of peptides identified across all search engines. A summary of identification counts and overlaps across search engines can be found in Supplementary Figure 1. The data from runs across all search engines is available at Mendeley [27].

To further assess the robustness of the model, we evaluated its performance on datasets not included in our training set and distinct from our Orbitrap, label-free, bulk, non-modified training data (Supplementary Table 2). We compared Tesorai Search against FragPipe (MSBooster), selected for its strong performance and ease-of-use in initial benchmarks. We compared the MS2 scan identification rates at a 1% FDR (Fig. 3). On a TMT-labeled dataset (Gabriel et al.) [28], we identified 13% more PSMs. On Time-of-Flight (TOF) instrument data, gains were +10% (Meier et al.) [29] and +30% (Van PuyVelde et al.) (Bruker timsTOF Pro), and +9% (Van PuyVelde et al.) (Sciex TripleTOF 6600+) [30]. Despite lacking specific PTM enrichment in training, Tesorai Search identified +51% PSMs on a phospho-enriched sample (Giasanti et al.). Furthermore, on challenging single-cell DISCO datasets (Lamanna et al. [31], 1 and 5 cell inputs), which differ significantly from bulk samples, we achieved 43-50% higher PSM identification rates. These results demonstrate that Tesorai Search generalizes effectively to diverse instruments, TMT labeling, PTM enrichment, and low-input single-cell data absent from its training.

**Figure 3:**
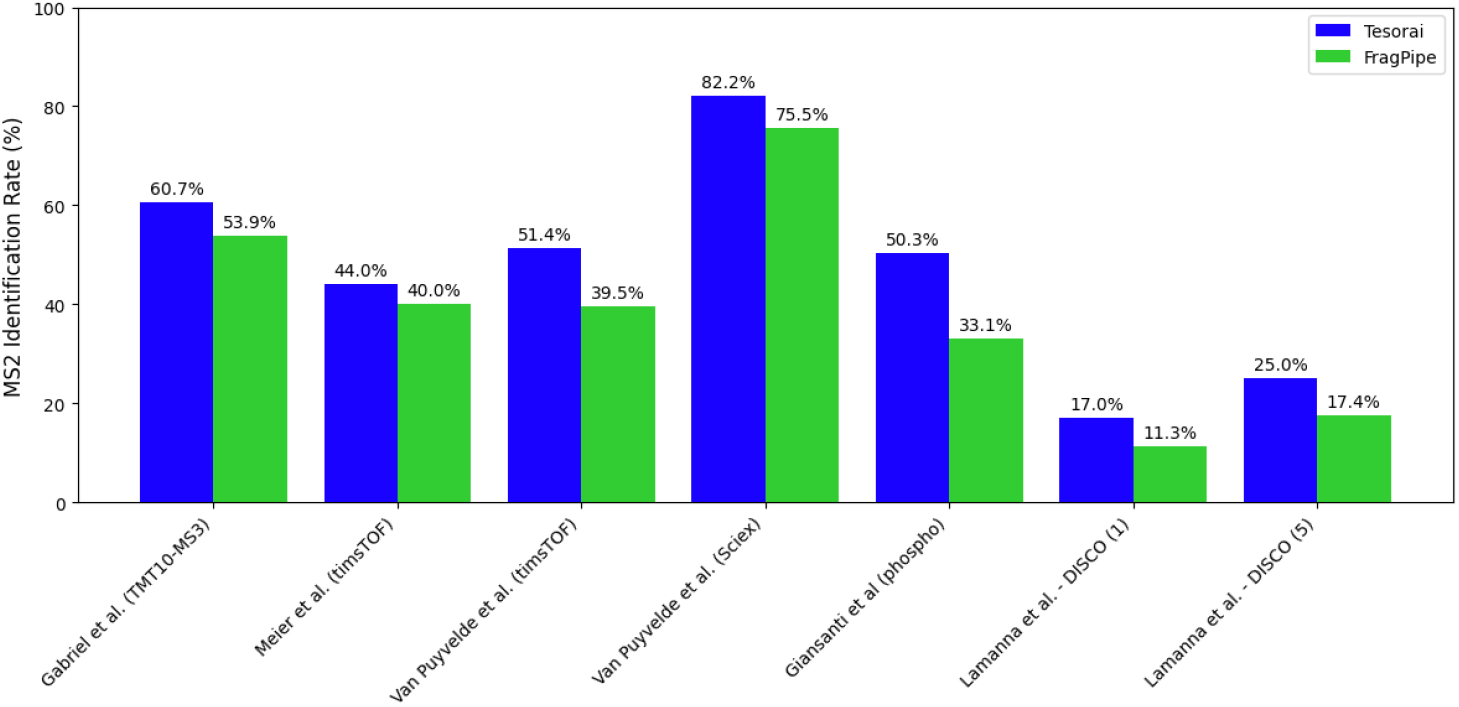
Scan (MS2) identification rate on data not seen in training. Comparison of MS2 scan identification rates at 1% FDR between Tesorai Search and FragPipe (MSBooster) across diverse proteomics datasets not seen during training: TMT10-MS3 (Gabriel et al.), TimsTOF instrument data (Meier et al.; Van Puyvelde et al.), Sciex data (Van Puyvelde et al.), phospho-enriched (Giasanti et al.), and single-cell DISCO (1 and 5 cells) (Lamanna et al.).

### Accurate false-discovery rate estimates

Increased identifications are meaningful only if the false-discovery rate (FDR) is correctly controlled, ensuring confidence in the results. We validated FDR control by measuring the false-discovery proportion (FDP), i.e. the true fraction of incorrect PSM or peptide matches, for user-defined FDR thresholds ranging from 0.1% to 10% (Fig. 4), ensuring FDP remained below the target FDR. Following [18, 32], we used entrapment analysis with the ISB18 dataset [33] (48 known target proteins: 18 synthesized, 30 contaminants) searched against a database combining these targets with the Ricinus communis (castor oil plant) proteome as an entrapment set chosen for low peptide overlap. PSM (Fig. 4A) and peptide (Fig. 4B) FDR were estimated using standard target-decoy competition. Given the high 668:1 entrapment-to-target peptide ratio, the FDP was computed as the number of identified entrapment sequences relative to non-entrapment sequences (known targets), using the conservative “combined” formula [34]. Our results demonstrate that the fraction of erroneous PSMs and peptides identified by Tesorai Search is consistently lower than the user-defined FDR threshold, confirming accurate control.

**Figure 4.**
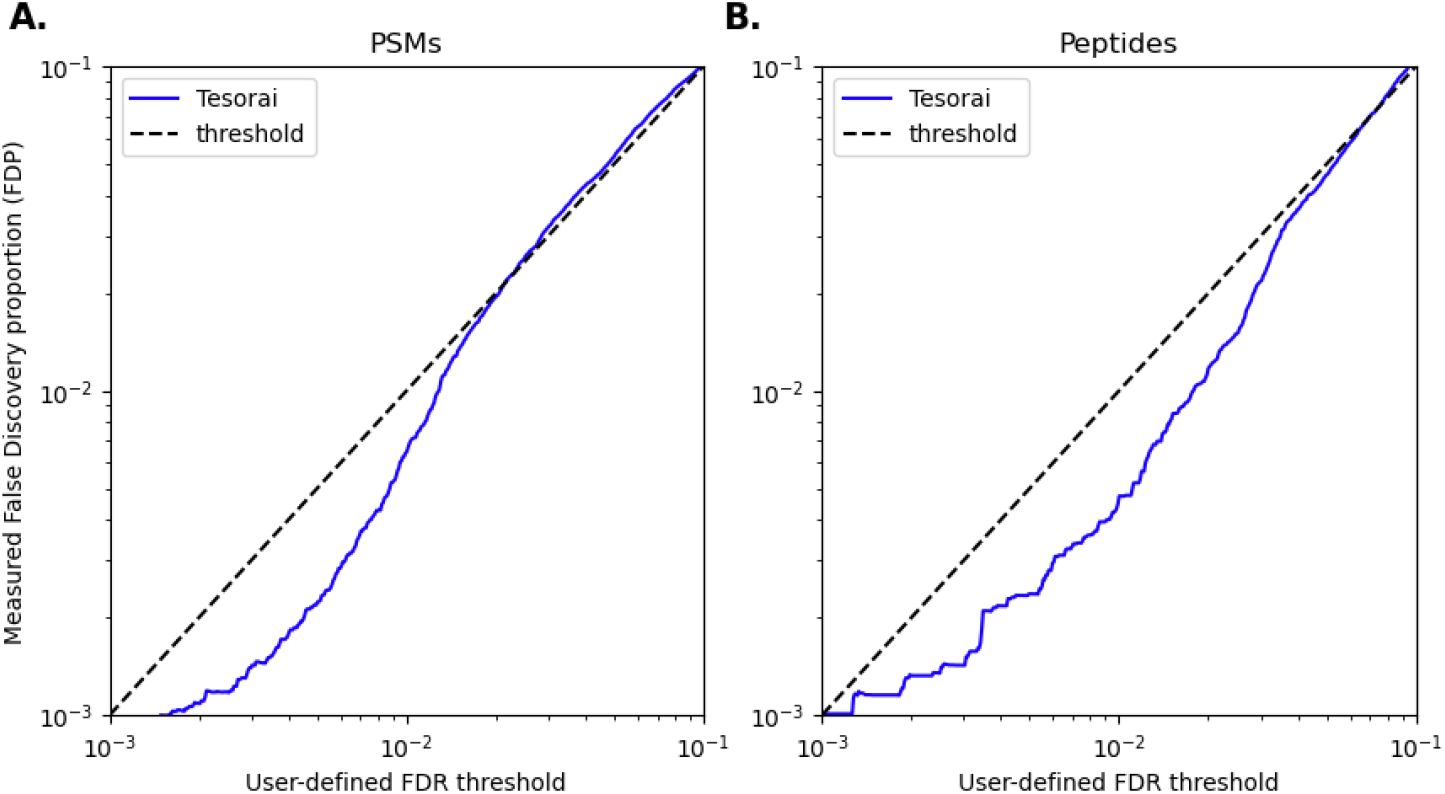
Measured false-discovery proportion (FDP) vs user-defined FDR threshold. **A. FDP vs FDR** at the PSM level of Tesorai Search (blue line), obtained through the entrapment method, compared to the user-defined false discovery threshold estimated using target-decoy competition in the region of interest, 0.1% to 10%. The dashed line represents the upper-bound case where the FDP is the same as the FDR. **B**. FDP vs FDR at the peptide level following the same evaluation approach.

### Newly identified peptides

Having established accurate FDR control, we next characterized the novel peptide identifications by investigating the score distributions and whether new peptides are corroborated by other search engines. Figure 5A-B, shows score distributions for decoys (orange) and targets (blue) on the Mel15 sample [24]. Target scores follow a bimodal distribution (Fig. 5A-B, horizontal marginal distribution) where the first mode closely matches the distribution of decoy PSMs. This bimodal distribution, which is more pronounced than for the Andromeda score (Fig. 5A, vertical marginal) and MSFragger Hyperscore (Fig. 5B, vertical marginal), enables cleaner score separation and makes the resulting identifications less sensitive to the FDR cutoff threshold. Most new peptides are also identified by MaxQuant or MSFragger, albeit at a 5% FDR cutoff (Fig. 5C-D).

**Figure 5.**
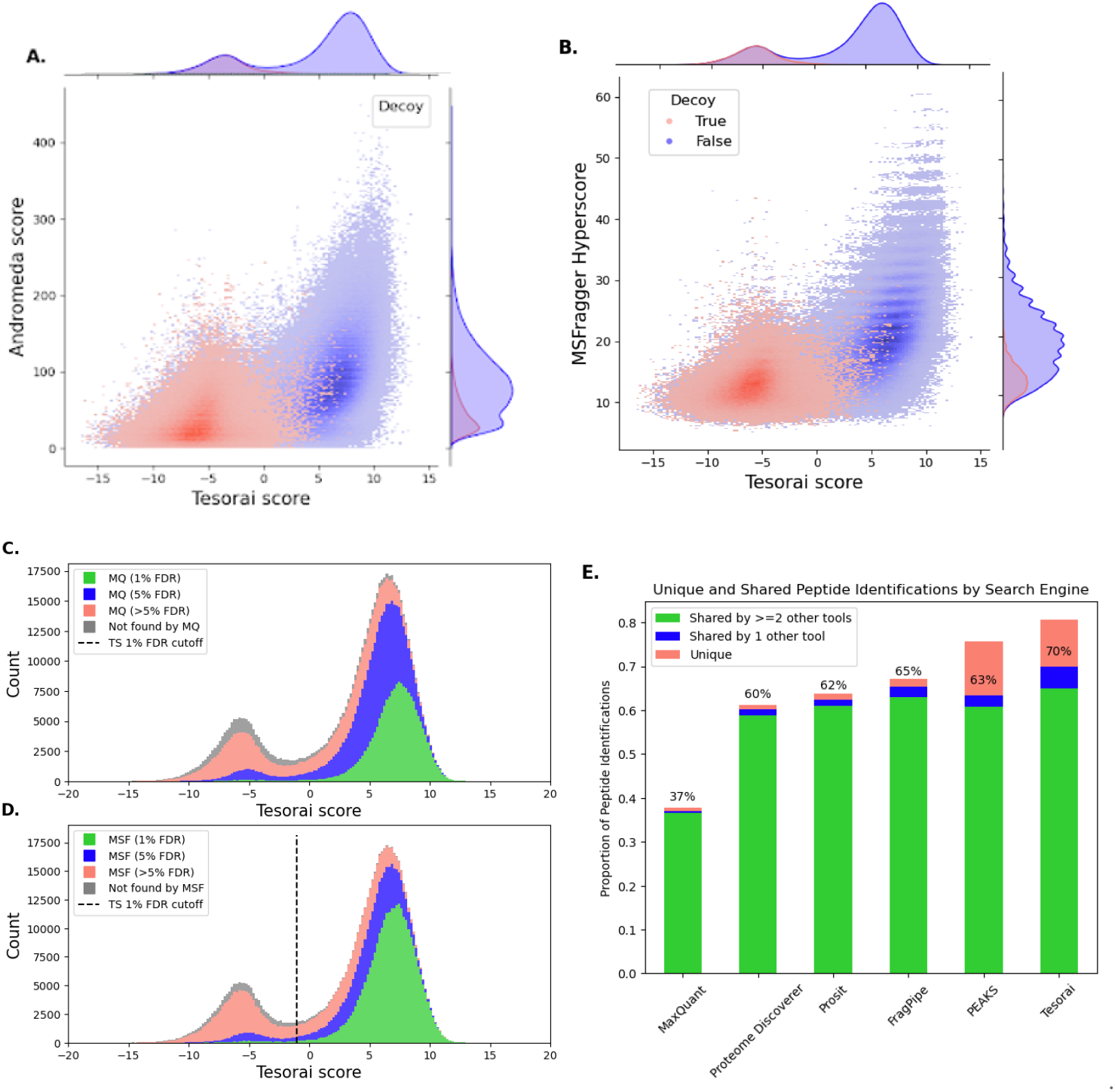
Newly identified peptides by origin using Bassani-Sternberg et al. (Mel15) dataset. **A-B**. Joint distribution of the Tesorai score against either the Andromeda score (A.) from MaxQuant or the MSFragger Hyperscore (B.). **C-D**. Breakdown of Tesorai Search identified PSMs by whether they were also identified by MaxQuant (C.) and MSFragger (D.) at various FDR thresholds. The dashed line represents the 1% FDR cutoff for the Tesorai Score. **E**. Breakdown of identified peptides from each tool, stratified by the number of other search engines that could confirm the identification. For Tesorai, we only included Proteome Discoverer and PEAKS in the tools used to confirm identification. We indicated the percentage of identified peptides that are shared with at least one other tool, as a percent of all peptides identified.

To further assess model reliability, we compared our peptide identifications against MaxQuant, Prosit, FragPipe, PEAKS, and Proteome Discoverer (Fig. 5E). For each tool, we categorized its identified peptides based on whether they were unique to that tool, shared with only one other tool, or shared with two or more other tools in the comparison set. For assessing our identifications, we did not count identifications that were only confirmed by FragPipe and MaxQuant, as these two tools are used by Tesorai Search to select initial PSMs for rescoring. Tesorai Search demonstrated the highest number of peptides confirmed by one or more other tools (70% of all identified peptides), compared to 37-65% for other tested tools.

### Model interpretation

To understand the contributions of different input features and internal representations to the model’s performance, we conducted an ablation study by corrupting model inputs and observing performance impacts (Fig. 6) On the PSM binary classification task used for model training, (validation set, ∼41% positive, see Methods), the uncorrupted model achieved 99.5% accuracy. As a baseline representing complete information loss, shuffling both the peptide sequence input and the associated theoretical fragment m/z values reduced accuracy to 58.6%, equivalent to random guessing.

**Figure 6.**
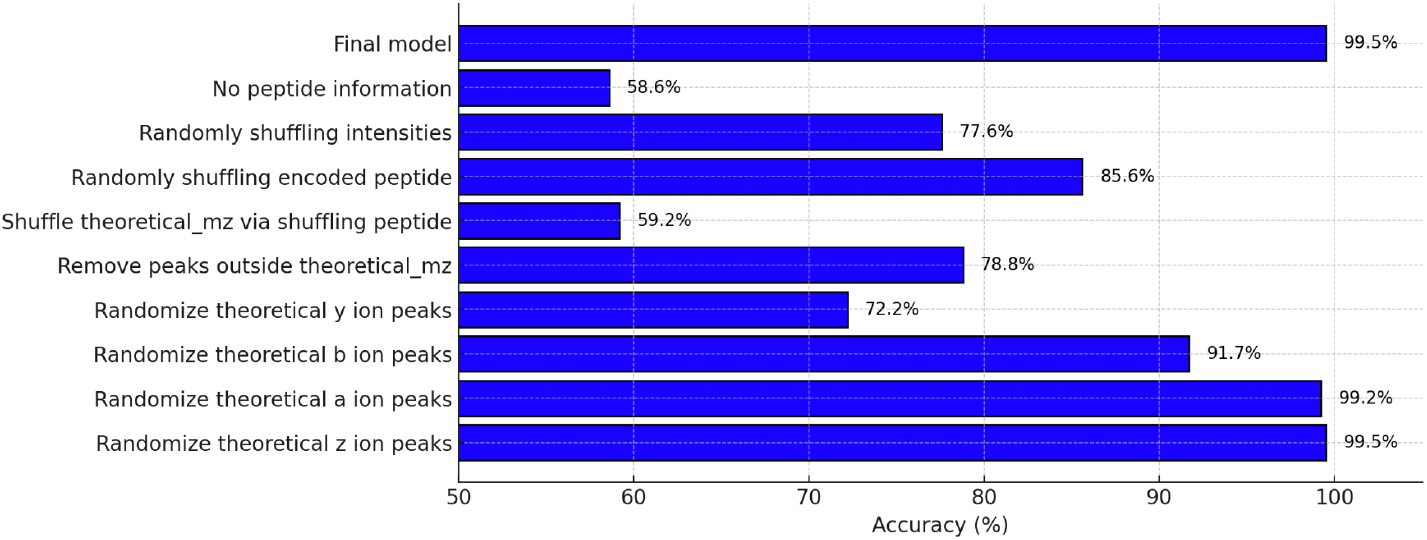
Effect of corrupting data inputs on model performance. Model accuracy (%) on the validation subset (41.4% positive, 58.6% negative) for the PSM binary classification task. Accuracy is shown for the full model (99.5%) and after selectively corrupting inputs via shuffling or randomization. No peptide information corresponds to shuffling all inputs.

We assessed input importance by selectively randomizing features. Randomizing experimental spectrum intensities (keeping m/z fixed) substantially dropped accuracy to 77.6%, showing that the model uses intensity information beyond m/z matching. Filtering the spectrum to only expected canonical fragment ion regions (a, b, y, z) also significantly reduced accuracy (78.8%), indicating that the model utilizes information from non-canonical peaks or their absence.

Investigating the peptide-derived inputs revealed a strong dependence on the theoretical fragment masses. Randomizing these reduced the accuracy to 59.2% (near baseline). Shuffling only the separate one-hot peptide sequence encoding had a lesser impact (85.6% accuracy), highlighting the importance of theoretical fragment matching over abstract sequence encoding. Examining individual theoretical ion types revealed that randomizing y-ions caused the most degradation (72.2% accuracy), followed by b-ions (91.7%). Randomizing a-ions (99.2%) or z-ions (99.5%) had minimal effect, aligning with expected CID fragmentation importance. Full results are in Supplementary Table 4.

### Cloud-based implementation allows for fast processing

To enhance processing speed and accessibility of proteomic analyses, we implemented a cloud-based interface for Tesorai Search. Users can perform database searches directly through a web browser without requiring local computational resources or software installations. We were able to re-process 250 immunopeptidomics samples from PXD019643 [35] in under 45 minutes, demonstrating the scalability of our implementation even on challenging workloads. This cloud-native approach not only simplifies workflows but also ensures that cutting-edge computational resources are available to researchers globally, facilitating rapid data processing and timely generation of results.

## Discussion

This paper introduces Tesorai Search, an end-to-end deep learning model that fundamentally shifts the paradigm for peptide-spectrum matching in mass spectrometry proteomics. Our core hypothesis is that the full spectrum and peptide sequence contain sufficient information for top-tier peptide identification, challenging the need for intricate feature engineering or auxiliary predictive models (like retention time or fragment intensity prediction) used in recent tools. Our model directly learns complex, non-linear associations between complete MS/MS spectra and peptide sequences, demonstrating, through extensive benchmarking, that this approach surpasses both traditional and machine learning-enhanced search engines.

Existing machine learning-enhanced search engines often combine dozens of handcrafted or predicted features, frequently using post-processing classifiers like Percolator trained on-the-fly with target-decoy strategies. While improving upon traditional methods, this added complexity can introduce variability and potential inaccuracies, particularly concerning FDR estimation. The fundamental issue [18] is that Percolator uses decoys for two distinct purposes: training a machine learning model to differentiate between correct and incorrect identifications and estimating the FDR. A key conceptual advance of Tesorai Search is its ability to learn directly from real peptide-spectrum matches (PSMs) without relying on decoys for model training. Trained once on a large-scale dataset of ∼289 million high-confidence, real PSMs, the model learns the genuine characteristics of correct identifications without potential biases introduced by artificial decoy sequences.

This results in a large increase in identification depth, with considerably more peptides identified than leading platforms like FragPipe, PEAKS, and Proteome Discoverer (12%, 9%, and 21%, respectively) across various use cases, including challenging immunopeptidomics and low-input samples. This enhanced sensitivity holds the potential for deeper biological insights [36]. For instance, in immunopeptidomics, identifying a broader repertoire of MHC-presented peptides is crucial for advancing cancer immunotherapy and vaccine design. The high proportion of identifications corroborated by other search engines supports their reliability. We also observed improved score separation compared to metrics like Andromeda, potentially due to better score calibration across different scans resulting from the classification-based training objective.

Furthermore, the model exhibits remarkable robustness and generalizability. Though trained exclusively on Orbitrap data, it performs well (identifying up to 50% more MS2 scans than FragPipe) on diverse sample types and data from instruments not included in the training set (TOF instruments, TMT-labeled samples). The end-to-end architecture implicitly learns necessary data processing steps, simplifying the workflow by obviating the need for explicit spectrum deisotoping, mass calibration, or charge correction. This inherent simplicity, combined with its implementation as a scalable, cloud-native platform, makes Tesorai Search readily accessible, facilitating rapid analysis of massive datasets and democratizing high-performance proteomics.

Several areas for future research emerge from this study. Though we focused exclusively on data-dependent acquisition workflows, the fundamental idea of using a single pretrained model for PSM rescoring is likely transferable to data-independent acquisition, promising simpler workflows, enhanced peptide identification, and improved FDR control in that setting as well. Although our method achieves state-of-the-art performance in the number of identified phosphopeptides, accurately localizing PTMs remains a challenge in the field. Furthermore, despite the advancements presented, overall scan identification rates can still be improved; combining our deep learning strategy with established techniques like match-between-runs and open window search approaches represents a promising direction for future research to maximize peptide recovery. Importantly, even with the strong concordance observed with existing tools, rigorous biological validation of uniquely identified peptides within relevant experimental contexts is essential to confirm their functional significance.

In conclusion, Tesorai Search demonstrates that a pre-trained deep learning model focused purely on the spectrum-sequence relationship can outperform other modern search engines. By offering enhanced sensitivity, robust FDR control, simplicity, and accessibility, this approach promises to accelerate discovery in fundamental biology and clinical proteomics.

## Methods

### Training dataset

For training, we identified 21 publicly available datasets (Supplementary Table 1), all processed with Orbitrap (Thermo Fisher Scientific) instruments. We first parsed the .raw files with ThermoRawFileparser to assemble the list of MS2 spectra. Each spectrum was represented as a tuple of two lists of floats, of equal lengths, representing the m/z value and the intensity of each (centroided) peak. No mass calibration or de-isotoping step was taken on the raw spectra.

In case the dataset was already processed with MaxQuant and an msms.txt file was publicly available, we used it directly, otherwise, we reprocessed the dataset with MaxQuant ourselves. Additionally, we processed some of the datasets with Comet and MS-GF+. Altogether, this resulted in a total of 289 million PSMs in the training dataset. We stored these spectra, the associated peptide sequences, and labels as TFRecords on disk to ensure low read times and high GPU utilization. To ensure that only high-quality PSMs were included in the training dataset, we filtered out those with an Andromeda score lower than 25, those with comet_score greater than 1e-3, and those with MS-GF+ SpecEValue greater than 1e-10.

We randomly split the peptide sequences based on mass into three splits: train, val, and test. Each peptide-spectrum match was then assigned to one of the buckets accordingly. Furthermore, we also reserved a set of PRIDE datasets where none of the PSMs were included in the train set. Finally, we created a list of benchmark datasets (see section “Processing of external data”), none of which were included in the training set, to measure the changes in identification rates.

### Model architecture

**Spectrum processing** happens on the CPU. We take as input a float32 vector of m/z values and a float32 vector of intensities. Spectra with fewer than 10 peaks are discarded. For spectra with more than 1100 peaks, only the 1100 most intense peaks are kept.

**Sequence processing** takes as input an int32 vector of one-hot-encoded amino-acid sequence of length s. It then computes theoretical fragment ions for a, b, y, z ions, two possible charge states (1 and 2), and optional water loss and ammonia loss. This results in 24 potential ions for each breakpoint in the peptide sequence. We used a total of 32 tokens to encode amino acids, where, in addition to the 20 canonical ones, we encoded the following 12 common modifications when computing theoretical fragment ions:

- M(ox) for methionine oxidation, (mass=147.035405 Da)
- Q(de) and N(de) for deamidation (mass += 0.9840 Da)
- Z for n-term acetylation (mass = 42.0106 Da)
- K(ac) for lysine acetylation (mass += 42.0106 Da)
- E(py) for pyroglutamic acid (mass -= 18.01057 Da)
- Q(py) for pyroglutamic acid (mass -= 17.02655 Da)
- S(ph), T(ph) and Y(ph) for phosphorylation (mass += 79.966331 Da)
- (tmt) and K(tmt) for isobaric labelling (variable mass shift depending on the --plex number) Furthermore, we trained our models with a fixed cysteine carbamidomethylation (+ 57.02146 Da).

### Loss function and model training

We used AdamW [40] for gradient descent, `torch.distributed` for distributed data-parallel multi-GPU training, and `torch.cuda.amp.autocast` for automatic mixed-precision training on float16 weights, to speed up the training process.

The model was trained on a single node with 8 Nvidia Tesla V100-SXM2-16GB GPUs, with a batch size of 512 and 1M steps, which corresponds to two full epochs over the training dataset and took a total of 64 hours. We used the PyTorch framework [37].

### Inference

For inference, the model takes as input a (sequence, spectrum) pair, and processes it in the same way as the training data. The spectrum is centroided and represented as a float32 vector of m/z values and a float32 vector of intensities. Spectra with fewer than 10 peaks are discarded. For spectra with more than 1100 peaks, only the 1100 most intense peaks are kept.

The sequence is one-hot encoded, and theoretical fragment ions are computed. It is then processed into the sequence encoder module, combined with the spectrum vector, and processed through the PSM score module.

We take the final number from the PSM score module as the score for the putative PSM, without passing it through the sigmoid function.

### Search engine configurations

#### FragPipe configuration

FragPipe analysis was performed within the FragPipe v22.0 computational platform. FragPipe includes MSBooster, Percolator, and Philosopher [41] tools by default for the downstream processing of the MSFragger search results. Decoys were generated using Philosopher, within the FragPipe computational platform. The provided TMT10-MS3 FragPipe workflow setting was used for the Gabriel et al. dataset. FragPipe search was performed without contaminants. The analysis included variable modifications for methionine oxidation, N-terminal acetylation, and fixed cysteine carbamidomethylation. Dataset-specific parameters are described for each dataset separately. All other parameters were kept default across all the experiments.

#### Tesorai Search configuration

Searches using Tesorai were performed with the following default settings unless noted otherwise. The default enzyme selected was Trypsin/P (`enzyme`: Trypsin/P), configured for specific cleavage (`enzyme_mode`: SPECIFIC). By default, the search didn’t include common contaminant sequences (`include_contaminants`: FALSE). Peptide identifications were filtered based on a default FDR threshold of 1% (`fdr_threshold`: 0.01). The minimum considered peptide length was set to 7 amino acids (`min_peptide_length`: 7). The maximum peptide length depended on the enzyme mode: for the default specific digestion, it was 63 amino acids (`max_peptide_length`: 63), while for unspecific digestion, the default maximum length was 25 amino acids. Default variable modifications are Methionine Oxidation and N-terminal Acetylation (`variable_modifications:` OXIDATION_M,ACETYLATION_NTERM). Cysteine Carbamidometylation is also enabled by default (`static_modifications:` CARBAMIDOMETHYL_C).

#### PEAKS configuration

PEAKS v12.5 was used with default settings. De novo search was disabled. Minimum and maximum peptide lengths were set to 7 and 50, respectively, except in immunopeptidomics runs, where we used 8 and 15.

#### Proteome Discoverer configuration

Proteome Discoverer v3.0 was used, and Chimerys rescoring was enabled, except in immunopeptidomics samples where it is not supported and on which Inferys was used. Minimum and maximum peptide lengths were set to 7 and 50, respectively, except in immunopeptidomics runs, where we used 8 and 15.

### Processing of external data

#### A. Bekker-Jensen (HeLa) tryptic dataset

Following [7], the Bekker-Jensen et al. [18] multi-protease dataset was downloaded from the PRIDE repository with the identifier PXD004452. Files mapping to the identifier `QE3_UPLC9_DBJ_SA_46fractions` were selected for analysis. Prosit results were obtained at PXD010871, in the zipped folder `Figure_2_3_5_Multiprotease_Dataset`, under the directory: `/trypsin/percolator_unzipped/prosit_target.psms`. MaxQuant results were obtained at PXD004452, under the directory: `SearchResults/msms.txt`.

The default settings were used for Tesorai search, except that the peptide length range was set to 7 to 50. All other settings were left to their default values. A human Swiss-Prot protein sequence database including annotated isoforms (downloaded May 3rd, 2024; 42421 protein sequences) was used for processing.

FragPipe results were run by us using the settings described above. Enzymatic digestion was set to `stricttrypsin,` and the peptide length range was set to 7 - 50.

#### B. Bassani-Sternberg et al. (HLA I immunopeptides) dataset

Following Wilhelm et al. [42], the HLA Class I sample from patient Mel15 [Bassani-Sternberg et al.] was downloaded from the PRIDE repository with the identifier PXD004894. Only files mapping to HLA Class I from patient Mel15 were used in the analysis. The sequence database (`HUMAN_2014`) was provided within the zipped `Search` folder in PXD004894, which was used in all experiments.

MaxQuant and Prosit results were obtained from PXD021398. MaxQuant results (`msms.txt`) were obtained from the zipped folder `Figure_5_Mel15_MaxQuant1`. Prosit results (`prosit_target.peptides`) were obtained from the zipped folder `Figure_5_Mel15_MaxQuant100_and_Rescoring`, under the directory `forPride_100%/rescoring_for_paper_2/percolator`.

Tesorai search enzyme mode was set to `UNSPECIFIC` with the minimum peptide length set to 8 and maximum to 15, following [42].

FragPipe results were run by us using the settings described above. Enzymatic digestion was set to `NONSPECIFIC`. The minimum peptide length was set to 8 and the maximum to 15.

#### C. Nowatzky et al. (HLA I immunopeptides) dataset

The Nowatzky et al. dataset was downloaded from MSV000089312. Only files mapping to identifiers `190514_H_FreyaLC_AD_Nowatzky_HLA_test` (KO and WT) were used in the analysis.

Tesorai search enzyme mode was set to `UNSPECIFIC` with the minimum peptide length set to 7 and the maximum to 15.

FragPipe results were run by us using the settings described above. Enzymatic digestion was set to `NONSPECIFIC`. The minimum peptide length was set to 7 and the maximum to 12.

Search was performed with a combined FASTA of Human and a few select pathogen proteomes (Uniprotkb, accessed 2023_12_07).

#### D. Meier et al. (HeLa sample, timsTOF)

Human cervical cancer cell (HeLa) dataset used in [24] was downloaded from the PRIDE repository with the identifier PXD010012. The 200ng, 100ms experiment was reproduced with the raw files provided in the zipped folder `HeLa_200ng_100ms_raw`. MaxQuant results (`msms.txt`) were obtained in the same PRIDE repository in the zipped folder `HeLa_200ng_100ms_txt`.

The default settings were used for Tesorai search.A reference human proteome (UP000005640) was downloaded from UniProt using one protein sequence per gene (March 26th, 2024; 20,590 protein sequences).

FragPipe results were run by us using the settings described above. Enzymatic digestion was set to `stricttrypsin` and the peptide length range was set to 7 - 63.

#### E. Gabriel et al. (HeLa-Yeast, TMT)

Human Yeast Dilution TMT labeled dataset [23] was downloaded from the PRIDE repository with the identifier PXD030340. The HCD fragmentation on the Orbitrap mass analyzer (HCD/OT) experiment was reproduced. The `190416_FPTMT_MS3_HCDOT_R1` raw file was used in the analysis. MaxQuant (`msms.txt`) and MaxQuant with Andromeda (`andromeda_target.pepides`) results were obtained in the same PRIDE repository in the zipped folder `HCD_OT_Without_rescoring`. Prosit rescored results (`prosit_target.peptides`) were obtained in the same PRIDE repository in the zipped folder `HCD_OT_With_rescoring`.

The default settings were used for Tesorai search, except the peptide length range was set to 7 to 50. A reference human proteome (UP000005640) was downloaded from UniProt using one protein sequence per gene (March 26th, 2024; 20,590 protein sequences). A reference Baker’s yeast proteome (UP000002311) was downloaded from UniProt using one protein sequence per gene (March 28th, 2024; 6,060 protein sequences).

MSFragger results were run by us using the settings described above. Enzymatic digestion was set to `stricttrypsin`. The provided `TMT10-MS3` FragPipe workflow setting was used and the peptide length range was set to 7 - 50.

#### F. Williams et al. (Single-cell NanoPOTS)

Following Yang et al. [7], the single-cell data from the nanoPOTS platform [21] was downloaded from the MassIVE repository under the identifier MSV000085230. We reproduced the experiment from using 1, 3, 10, and 50 cells from the MCF10A 30-minute experiment. We used the human Swiss-Prot protein sequence database provided at MSV000085230 for all the experiments (`UniprotKB_homosapiens_Swiss_Prot_122916`).

The default settings were used for Tesorai search. FragPipe results were run by us using the settings described above. Enzymatic digestion was set to `stricttrypsin` and the peptide length range was set to 7 - 63.

#### G. Van Puyvelde et al. (BrukertimsTOF, Sciex)

Thermo Orbitrap QE HF-X, Bruker timsTOF Pro, and Sciex TripleTOF 6600+ [25] data was downloaded from the PRIDE repository with the identifier PXD028735. Raw files mapping to `LFQ_[timsTOFPro_PASEF, TTOF6600]_DDA_Condition_A_Sample_[Alpha, Beta, Gamma]_[01-04]`were selected for the analysis. Sciex .wiff files were peak-picked with MSConvert [43] and converted to mzML file format as our current platform does not natively support .wiff format. A reference database containing the Human, Yeast, and E.coli protein sequences was downloaded from UniProt in May 2024.

The default Tesorai search settings were used, except the peptide length range was set to 7 - 50. FragPipe results were run by us using the settings described above. Enzymatic digestion was set to `stricttrypsin` and the peptide length range was set to 7 - 50.

#### H. Lamanna et al. (Single-cell DISCO)

Following [8], the single-cell data from the DISCO platform [26] was downloaded from the PRIDE repository under the identifier PXD019958. We reproduced the experiment from [8] using 1 and 5 cells Orbitrap Thermo Q-Exactive HF-X raw files, using all the 3 replicas for each experiment. We used the human Swiss-Prot protein sequence database provided at MSV000085230 for all the experiments (`UniprotKB_homosapiens_Swiss_Prot_122916`) that was also used in the Single-cell NanoPOTS experiment.

The default Tesorai search settings were used.FragPipe results were run by us using the settings described above. Enzymatic digestion was set to `stricttrypsin` and the peptide length range was set to 7 - 63.

#### I. Giansanti et al. (phosphopeptidomics) dataset

The following six Orbitrap files were downloaded from the PRIDE repository under the identifier PXD001428 from Giansanti et al. [44]:

- OR8_130622_TT_Trypsin_Ti-IMAC_Rep1_B1.raw
- OR8_130622_TT_Trypsin_Ti-IMAC_Rep1_B2.raw
- OR9_20130628_TT_Trypsin_Batch2_R1.raw
- OR9_20130628_TT_Trypsin_Batch2_R2.raw
- OR9_20130628_TT_Trypsin_Batch3_R1.raw
- OR9_20130628_TT_Trypsin_Batch3_R2.raw

The default settings were used for Tesorai search with an additional variable modifications enabled for phosphorylation modification of serine (S), threonine (T), and tyrosine (Y) residues (PHOSPHORYLATION_STY) and peptide length range between 7 and 50.

FragPipe was run with the default settings and phosphorylation STY variable modification enabled in the MSFragger search parameters. Peptide length range was between 7 and 50.

### Cloud-based implementation

We enabled fast and scalable processing by running with high parallelism and leveraging on-demand, serverless Cloud resources. By default, each sample file is first run through a combination of FragPipe (MSFragger), MaxQuant and Comet at 100% FDR to generate a list of candidate PSMs for each MS2 scan. For immunopeptidomics runs, we disable MaxQuant and Comet to speed-up processing. This comes with almost no reduction in performance. Data collected on Sciex (.wiff) or Bruker (.d) instruments also is processed by MSFragger only.

Every file and algorithm is run on a separate Cloud machine, with optimized resources for that stage. The pre-trained model runs on T4 Nvidia GPUs and rescores the candidate PSMs.

These rescored PSMs are then filtered at 1% FDR, and assembled into peptide and protein lists. We use an open-source FFIA [45] quantification algorithm for all Orbitrap, Sciex and mzML datasets. For Bruker, we leverage the default quantification in FragPipe (IonQuant) [46].

The final results are surfaced in an intuitive and easy-to-use web interface, and full psm, peptide and protein tables are downloadable directly from the platform.

### FDR estimation

We estimate FDR with the standard target-decoy approach. We use the default settings of each search engine to generate decoy sequences from the user-provided FASTA file. For MaxQuant and FragPipe, this is done by reversing the protein sequences prior to in-silico digestion. For Comet, we kept default settings: Comet generates decoys by reversing each target peptide sequence, keeping the N-terminal or C-terminal amino acid in place. We do not use protein-level information for additional filtering (sequential FDR).

### Entrapment experiment to measure false-discovery proportion

The raw data for the entrapment analysis was accessed on March 15, 2024 at https://regis-web.systemsbiology.net/PublicDatasets/18_Mix/Mix_7/LTQ/RAW_Data/. Following [29], we analyzed files 2-10 and excluded number 11. The Castor plant proteome was accessed from Uniprot on 2024-04-25. The list of ISB18 proteins was downloaded from https://regis-web.systemsbiology.net/PublicDatasets/database/18mix.fasta.

Following [29], we used strict tryptic enzymatic in-silico digestion (this is our default), and set missed cleavages to 0 (default 2). To reduce the probability of a contaminant randomly matching with a peptide from the castor plant, we set the minimum peptide length to 8. All other settings were kept default. All peptide mappings accounted for Isoleucine to Leucine substitution.

## Data availability

All raw data analyzed in this study were already publicly available. All results run by ourselves, from Tesorai Search, MaxQuant, FragPipe, PEAKS, and Proteome Discoverer will be made available on Mendeley Data [22].

## Code availability

The Tesorai platform, used to generate the main results, is available online at console.tesorai.com. The code used to process the results and generate figures and tables is available at https://github.com/tesorai/tesorai_search.

## Footnotes

1 https://github.com/tesorai/tesorai_search

