## Supplementary Notes for "Back to Basics: Spectrum and Peptide Sequence are Sufficient for Top-tier Mass Spectrometry Proteomics Identification"

### Tesorai Search – Supplementary materials

#### 1. Training Dataset statistics

| PRIDE Project | Sample Count | Species | Sample Type |
|---|---|---|---|
| PXD024364 | 65349 | Homo sapiens (human) | Cell |
| PXD010154 | 53374 | Homo sapiens (human) | Tissue |
| PXD021013 | 29284 | Homo sapiens (human) | Synthetic |
| PXD003668 | 24565 | Homo sapiens (human) | Cell |
| PXD013615 | 19756 | Homo sapiens (human) | Cell |
| PXD004732 | 16264 | Homo sapiens (human) | Synthetic |
| PXD037285 | 16035 | Homo sapiens (human) | Cell |
| PXD014877 | 13426 | Multiple* | Not sure |
| PXD023119 | 10835 | Homo sapiens (human) | Synthetic |
| PXD010595 | 10602 | Homo sapiens (human) | Synthetic |
| PXD005353 | 6370 | Homo sapiens (human) | Cell |
| PXD019483 | 6133 | Homo sapiens (human) | Cell |
| PXD023120 | 5367 | Homo sapiens (human) | Synthetic |
| PXD020079 | 3260 | Homo sapiens (human) | Cell |
| PXD001608 | 2299 | Homo sapiens (human) | Tissue |
| PXD004977 | 1505 | Homo sapiens (human) | Cell |
| PXD000955 | 1347 | Saccharomyces cerevisiae (baker's yeast) | Cell |
| PXD001695 | 1231 | Saccharomyces cerevisiae (baker's yeast) | Cell |
| PXD008127 | 783 | Homo sapiens (human) | Tissue |
| PXD020011 | 660 | Homo sapiens (human) | Cell |
| PXD001865 | 434 | Saccharomyces cerevisiae (baker's yeast) | Cell |

| | | | |
|---|---|---|---|
| PXD002452 | 33 | Mus musculus (mouse) | Cell |

**Supplementary Table 1.** *Summary of PRIDE project data by species.* This table lists the sample counts for different PRIDE projects along with their associated species.

*PXD014877: Vitis vinifera (grape), Danio rerio (zebrafish) (brachydanio rerio), Escherichia coli, Rattus norvegicus (rat), Gallus gallus (chicken), Arabidopsis, thaliana (mouse-ear cress), Drosophila melanogaster (fruit fly), Homo sapiens (human), Caenorhabditis elegans, Triticum aestivum (wheat), Dictyostelium, discoideum (slime mold), Saccharomyces cerevisiae (baker's yeast), Bos taurus (bovine), Mus musculus (mouse), Sus scrofa domesticus (domestic pig)

#### 2. Model accuracy

| Metadata | Bucket | Sample Count | Accuracy |
|---|---|---|---|
| Peptide Length | [7, 10] | 43346 | 98.9 |
| Peptide Length | (10, 12] | 27766 | 99.4 |
| Peptide Length | (12, 14] | 16833 | 99.8 |
| Peptide Length | (14, 16] | 18978 | 99.7 |
| Peptide Length | (16, 18] | 17081 | 99.8 |
| Peptide Length | (18, 20] | 16379 | 99.9 |
| Peptide Length | (20, 22] | 6617 | 99.8 |
| Peptide Length | (22, 24] | 3970 | 99.9 |
| Peptide Length | (24, 50] | 3980 | 99.7 |
| MZ ratio | [0, 500] | 21776 | 99.1 |
| MZ ratio | (500, 600] | 29199 | 99.1 |
| MZ ratio | (600, 700] | 32453 | 99.3 |
| MZ ratio | (700, 800] | 22204 | 99.8 |
| MZ ratio | (800, 900] | 15257 | 99.8 |
| MZ ratio | (900, 1000] | 16225 | 99.7 |
| MZ ratio | (1000, 1500] | 20918 | 99.9 |
| Fragmentation | CID | 35950 | 98.9 |
| Fragmentation | ETCID | 499 | 99.6 |

| Metadata | Bucket | Sample Count | Accuracy |
|---|---|---|---|
| Fragmentation | ETD | 3694 | 99.6 |
| Fragmentation | ETHCD | 2708 | 99.6 |
| Fragmentation | HCD | 115181 | 99.6 |
| Hardware | FTMS | 84914 | 99.5 |
| Hardware | ITMS | 73118 | 99.5 |
| Species | Arabidopsis thaliana | 134 | 98.5 |
| Species | Escherichia coli | 1819 | 99.9 |
| Species | Homo sapiens | 125886 | 99.4 |
| Tryptic | FALSE | 60855 | 99.5 |
| Tryptic | TRUE | 97177 | 99.5 |
| PTM present | FALSE | 147181 | 99.5 |
| PTM present | TRUE | 10851 | 99.3 |
| **TOTAL** | **All Data** | **158032** | **99.5** |

**Supplementary Table 2.** *Summary of model performance across different metadata.* The table presents the sample counts and model accuracy (%) for various metadata, including peptide length, MZ ratio, fragmentation type, hardware type, species, trypticity, and presence of post-translational modification (PTM). Fragmentation type, hardware, species, trypticity, and PTM are categorical features. Accuracy reflects the percentage of correct predictions made by the model within each feature group. The model shows consistent high performance across most feature groups, with the majority of accuracy values above 99%.

| Category | Dataset | FragPipe | Tesorai |
|---|---|---|---|
| TMT | Gabriel et al. (TMT10-MS3) | 6,291 | **6,748** |
| TOF instruments | Meier et al. (timsTOF) | 67,155 | **68,377** |
| | Van Puyvelde et al. (timsTOF) | 68,065 | **75,199** |
| | Van Puyvelde et al. (Sciex) | 34,809 | **40,886** |
| Phosphopeptidomics | Giansanti et al. | 12,588 | **14,387** |
| Single-cell(s) | 1 cell | 5,713 | **6,744** |

| | | | |
|---|---|---|---|
| Lamanna et al. - DISCO | 5 cells | 9,598 | **11,134** |
| Immunopeptidomics | Sarkizova et. al. | 11,841 | **12,063** |

**Supplementary Table 3.** FragPipe vs Tesorai

| Ablation experiment | Accuracy |
|---|---|
| **Final model** | **99.5** |
| No peptide information (Shuffle peptide and theoretical_mz) | 58.6 |
| Randomly shuffling intensities | 77.6 |
| Randomly shuffling encoded peptide | 85.6 |
| Shuffle theoretical_mz via shuffling peptide | 59.2 |
| Remove peaks in real spectrum outside of theoretical_mz fragment ions (a,b,y,z) | 78.8 |
| Randomize theoretical y ion peaks | 72.2 |
| Randomize theoretical b ion peaks | 91.7 |
| Randomize theoretical a ion peaks | 99.2 |
| Randomize theoretical z ion peaks | 99.5 |

**Supplementary Table 4.** Ablation experiment

| | MaxQuant | Proteome Discoverer | FragPipe | Peaks | Tesorai | union |
|---|---|---|---|---|---|---|
| bassani_sternberg | 22166 | 35882 | 39410 | 44396 | 47393 | 57935 |
| bekker_jensen | 169476 | 190362 | 174795 | 170604 | 183147 | 224741 |
| nowatzky | 3677 | 8333 | 10848 | 11255 | 11787 | 14107 |
| williams/1_cells | 1626 | 1964 | 1910 | 1892 | 2472 | 2761 |

| | | | | | | |
|---|---|---|---|---|---|---|
| williams/3_cells | 2758 | 3435 | 3611 | 4048 | 4301 | 5114 |
| williams/10_cells | 4488 | 5382 | 5620 | 6130 | 6614 | 7766 |
| williams/50_cells | 7890 | 10482 | 10720 | 12285 | 13030 | 15472 |

Supplementary Table 3. Top panel: Nowatzky. Middle panel: unknown. Bottom panel: Bassani-Sternberg

### Supplementary Methods

##### Sarkizova et al. (HLA I immunopeptides) dataset

The following five HLA-I files were downloaded from the MassIVE repository under the identifier MSV000084172 from Sarkizova et al. [31]:

- GG20161104_CRH_HLA_C0302_BioRep1_inject1.raw
- GG20161104_CRH_HLA_C1402_BioRep2_inject1.raw
- MS20160409_CRH_HLA_B_2705_BioRep1_TechRep1.raw
- YE_20180428_SK_HLA_A0202_3Ips_a50mio_R1_01.raw
- YE_20180517_SK_HLA_A1101_3IPs_a50mio_R1_01.raw

Tesorai search results were with contaminants turned off. The enzyme mode was set to `UNSPECIFIC` with the minimum peptide length set to 8 and maximum to 15.
FragPipe results were run by us using the settings described above. Enzymatic digestion was set to `NONSPECIFIC`. The minimum peptide length was set to 8 and the maximum to 15
